# The potential of parrotfish faeces in replenishing reefs with coral-associated microbiome

**DOI:** 10.1101/2020.09.15.298737

**Authors:** Trigal M. Velásquez-Rodríguez, Catalina Zuluaga-Arias, Sandra M. Montaño-Salazar, John M. González, Juan Armando Sánchez

## Abstract

*Sparisoma viride* is the most abundant parrotfish in the Caribbean and is considered as the most important excavator due to corallivore behaviour. Parrotfishes are a keystone group that favour the growth and resilience of coral reefs removing macroalgae and structuring the benthic communities. The microbial symbiotic communities are involved in multiple functions related to nutrition and immunity maintaining corals health. Because *S. viride* scrape coral tissues, the skeleton and the algae on coral, it could be important reservoirs or vectors of microorganism for the corals through the faeces dispersion, however, the role of parrotfishes as reservoirs are poorly studied. Establishing microbial communities present in parrotfish faeces will contribute to understand the ecological impact of parrotfishes in coral resilience. We investigated the composition of disseminated bacteria and the extent to which the cell integrity of dinoflagellate photosymbionts (Symbiodiniaceae) is maintained in the faeces and compare with sediments and water column controls. Then, we analysed diversity and structure of bacterial communities at family level and search similarities between faeces of the study and coral associated microbiome reported in the literature. Similar levels of structural integrity and photosynthetic health of Symbidiodinaceae cells were found in both faeces and reef sediments. Besides, the sediments microbiome echoes the parrotfish faecal microbiome by sharing high diversity and a similar bacterial community composition. Several bacterial families were present in parrotfish faeces and in coral microbiome reported in the literature highlighting the dispersal potential of parrotfishes replenishing coral reefs. Despite the sampling limitations, these findings uncover the potential role of the excavator parrotfish in enriching environmental reservoirs, especially reef sediments, with coral-associated bacteria and photosynthetic microalgae. Parrotfishes could reinforce the coral microbiome and facilitate coral symbiont acquisition, key features critical to maintaining the fitness of one of the most threatened ecosystems on Earth. This finding could be considered as a first step in uncovering a mechanism for reef-microbiome maintenance.

## Introduction

The coral microbiome includes species-specific symbiotic interactions with unicellular dinoflagellates of the family Symbiodiniaceae, endolithic green algae, bacteria, archaea, fungi, protists and viruses (Ricci et al. 2019). From the surface mucus layer into the skeleton, coral-associated bacterial communities and photosynthetic dinoflagellates support coral homeostasis (Ainsworth et al. 2015). Bacteria occupy specific niches in the coral according to their function at the intracellular and perialgal space (Hernandez-Agreda et al. 2017), where critical roles in catabolizing organic matter and recycling nutrients are performed (Bourne et al. 2016). Particularly, bacteria located in the coral mucus supply nutrition while protecting against pathogens and UV light, which is an emerging and dynamic process from the constant environmental interaction (Bourne et al. 2016). In addition, they produce diffusible and non-diffusible antibacterial compounds that, along with physical exclusion, regulate other bacterial populations influx (Shnit-Orland and Kushmaro 2009).

Acquisition of symbiotic dinoflagellates and bacteria by corals may occur mainly by horizontal transmission from the environment, occasionally by vertical transfer to the gametes from the parent colony, or by a mixture of both (Hernandez-Agreda et al. 2017). An essential element for the uptake process is the availability of microbial components outside the host (Littman et al. 2009), especially in environmental reservoirs such as the water column (Takabayashi et al. 2012), benthic macroalgae beds (Porto et al. 2008) and sediments (Nitschke et al. 2016). Pioneering observations found corallivorous fish delivering coral symbionts to the environment (Muller-Parker 1984). In the Caribbean, the occurrence of Symbiodinaceae cells in faeces of the stoplight parrotfish *Sparisoma viride*, with densities up to 38.3×10^3^ cells ml^−1^, has supported this notion (Porto et al. 2008; Castro-Sanguino and Sánchez 2012).

*S. viride* is the main excavating parrotfish of the Caribbean. This feeding behaviour leads to the ingestion of symbionts while foraging on rock and live coral (Sanchez et al. 2004; Francini-Filho et al. 2008). Notably, its primary substratum to feed is the epilithic algal matrix found on dead coral, and live coral is approximately 1% of the total fish diet (Bonaldo et al. 2014). It is important to remark, which is also well-documented in the literature, that *S. viride* targets only endolithic material for feeding (Bruggemann et al. 1994; Clements et al. 2016; Nicholson and Clements 2020). The ingestion of photoautotrophs, like Symbiodinaceae, should be considered as a by-product. Additionally, *S. viride* forages other benthic species such as sponges (Radax et al. 2012; Burkepile et al. 2019a). After excavating, the grinding action of *S. viride* pharyngeal jaws and long gut pass transforms calcium carbonate in fine sediment (Perry et al. 2015), which contains living Symbiodinaceae (Castro-Sanguino and Sánchez 2012) and even viable macroalgae (Vermeij et al. 2013).

Coral reefs face unprecedented rates of degradation and increasing stressful scenarios (Hughes et al. 2017). Uncovering processes and mechanisms to enhance ecological resilience is crucial in addressing this crisis (Chung et al. 2019). Moreover, abundance declining in key species such as *S. viride* (Near Threatened - IUCN Red List) is threatening the ecological input for local resilience. On this basis, we hypothesized that the dispersal of coral microbiome components is influenced by *S. viride* faeces suggesting a coral reef microbiome maintenance mechanism, which is important to maintain the environmental reservoirs of microbial communities that are key on coral reefs.

## Materials and Methods

### Data and code availability

The qiime2R and R code are available on the Github repository (https://github.com/smmontanos/Microbiome-analysis-of-Parrotfish.git) with a readme. The procedures are explained in the section of Methods Details.

Original raw sequences of the bacterial communities associated with the samples of the study were deposited in NCBI short read sequences (SRA) under the Bioproject PRJNA623459.

### Study Area

The fieldwork was carried out in the Archipelago of San Andrés, Old Providence, and Santa Catalina (Southwestern Caribbean), which is composed of oceanic islands, submerged banks, and atolls located approximately 800 km northwest of Colombia. This territory is part of the Seaflower Biosphere Reserve declared by UNESCO in 2005 and is considered the largest Marine Protected Area (MPA) in the Caribbean. Seaflower MPA is relevant at global scale as it contains over 200.000 ha of productive open-ocean reefs and extended reefs associated with diverse ecosystems (The CaMPAM Mapping Project, 2017), whereby it is regarded as the third largest barrier reef worldwide. Specifically, this study was developed around San Andrés Island which has 15 km of long barrier reef (The CaMPAM Mapping Project, 2017). The sampling was conducted at Nirvana Diving Site in 12° 30′ 05″N and 81° 43′ 56″ W and West Point reef at the site located in the coordinates 12°31’15.45”N, 81°43’48.60”W along the leeward reef of the island (“Fig. S1”). Sampling was possible thanks to the research permit N°00254 (2017) issued by Autoridad Nacional de Licencias Ambientales (ANLA), Ministerio de Medio Ambiente y Desarrollo Sostenible, Colombia

### Symbiodinaceae Faeces sampling

Observational samplings of focal male adult individuals (terminal phase) of *S. viride* were conducted using SCUBA diving in a total area of 5.000 m^2^, consisting of patch reefs (depth 3 to 12 m). The standardized time for focal observations was 10 minutes per individual; preferred individuals were those foraging near to the bottom due to the consistent quality of the faeces released. During this time, one pellet faeces sample per individual was collected immediately after released on substrate using 10 ml syringes, intensely procuring pellet material only. A total of 40 pellet faeces samples were collected. Additionally, sediment control (seabed) samples were taken (60 cm horizontally to the pellet faeces fall place). Each syringe was disposed in a Ziploc bag marked with the sample type and safely kept. After the immersion, samples were fixed in formalin 10 % to preserve cell integrity. This sampling was developed during June 2016. Underwater time corresponded to 24 diving hours.

### Faeces filtration

In the laboratory, each sample was filtered using a plastic Büchner funnel coupled with a vacuum pump and a previously weighted Whatman filter paper grade 4 (25-30um mesh size). Sample scattered on the filter paper was washed with 0.22 um Filtered Sea Water (FSW) until recovering a final volume of 50 ml. After, the filter was dried at room temperature, and its new weight with the sample sediment retained on it was registered. Symbiodinaceae cells size ranged from 5 to 20 um, and therefore, it was expected to recover them the filtrate. Each 50 ml falcon tube was centrifuged at 3100 rpm for 5 min. Subsequently, the pellet was resuspended into 250 ul of FSW. The difference between the blank filter paper and the dried sediment sample on filter paper was considered as the collected faeces’ weight dry sediment. This process was applied to the control samples as well. Filtered samples were protected from light incidence due to the possible autofluorescence degradation of Symbiodinaceae cells.

### Microscopy verification and enumeration

Microscopic observation to verify the presence of Symbiodinaceae cells was carried out on an Improved Neubauer haemocytometer (Hausser Scientific, USA), under a compound light microscope Olympus CX21, using 40X magnification. Symbiodinaceae recognition was based on previous observations of tissue samples and zooxanthellae extruded from *Pterogorgia guadalupensis*, and from published identification photos on scientific papers. Thus, cells were identified using characters as unicellular circular shape, golden-brownish colour, elongated and reticulated chloroplast and large pyrenoid body (MR. Nitschke, doctoral dissertation and personal communication, 2015).

### Viability quantification by flow cytometry

#### FCM Standardization

Quantification of viable cells (high fluorescence) and degrading/unviable cells (low fluorescence) of Symbiodinaceae in *S. viride* faeces was carried out by flow cytometry (FCM). First, it is important to note that viability was not based on fluorescence intensity only. Although Chl autofluorescence may prevent the separation between viable and apoptotic cells (Strychar et al. 2004), low Chl fluorescence is related to stressed or damaged Symbiodinaceae (Gierz et al. 2020), which cell morphology can be accurately spotted using FCM measurements emitted by particles digitalized as: first, thanks to the forward scatter (FSC) or the signal related to the cell size and shape; and second, due to side scatter (SSC) or the signal indicative of cell granularity. Based on FSC and FSC, parameters are specific to each particle type, FCM identifies cell populations composed of the same particle types. Moreover, FCM identifies cells populations based on: first, particles light scattering principles, and second the excitation and emission of fluorescence; both as the response to a light beam. In detail, a cell solution enters into the sample injection probe, in which there is a stable stream (generally PBS solution) that forces each particle to pass individually. During the travel, each particle passes through a flow cell, the site in which is stimulated using laser illumination (488 nm or 633 nm, BD FACs Canto II, BD). As a result, the particle emits a response expressed in two measures of light scattering that are detected and converted into electrical signals by a light detector.

For FCM standardization purposes, preliminary assays were developed using two different samples types; first *Pterogorgia guadalupensis*, a common zooxanthellate octocoral in the study area, extracts from live tissue, and second, *P. guadalupe*nsis zooxanthellae cultures aliquots. In the beginning, the process consisted in differentiating cellular components from debris particles based on FSC and SSC. Both parameters indicated heterogeneous size of cellular components in samples. In the next stage, cellular components were distinguished using fluorescence emission. For this, propidium iodide (PI) staining was performed to both types of samples as a way of promoting fluorescence resolution for cell viability quantifications; however, there were not observed significative differences in fluorescence, in comparison to unstained samples. Consequently, it was concluded that Symbiodinaceae autofluorescence signal was pertinent as a viability quantification criterion. In this way, a red fluorescence threshold set (670 nm long-pass), related to Symbiodinaceae autofluorescence was established after testing the efficiency of three different PMT (520 nm long-pass, 590 nm long-pass and 670 nm long-pass) in cells differentiation. Finally, cells populations in faeces and sediment samples were gated and counted using FCS, SSC and fluorescence parameters between particles. Subsequently, sorting (FACs ARIA II, BD) was carried out on the individual defined populations. After, each sorted population was observed using an Improved Neubauer haemocytometer and photographic images were taken. Additionally, fluorescence microscopy observations were done to corroborate light microscopy findings. Based on the sorted population results, the FCM settings were considered standardized for the *S.viride* faeces viability quantification.

#### FCM conditions for viability quantification

*S.viride* faeces samples and sediment controls filtered were analyzed using a BD FACS Canto II cytometer, coupled with the FACS Diva 6.1. Software. A sample volume of 200 ul was used for each run, and a total of 1500-2000 cell events were registered. FSC and SSC were collected using a 488nm band-pass filter, with blue light of origin. Additionally, Symbiodinaceae red autofluorescence signals (chlorophyll a) were collected via 670nm band-pass filter. As a conclusion, FSC and SSC, in combination with the Symbiodinaceae endogenous autofluorescence were used to discriminate zooxanthellae population from macroalgae fragments, non-algal cells and high quantity of debris in *S.viride* faeces. The same parameters functioned for the differentiation of Symbiodinaceae viable cells and degrading/unviable cells. Normalization of data was made for assuring an accurate comparison of cell size, fluorescence patterns and viability percentages between populations in the total samples. Details used for the FCM analysis, under logarithmic criterion, were (parameter/voltage): FSC/220, SSC/222, FITC/560, PE/500, PE/500 and PE-Cy7/600.

### Statistical Analysis for Symbiodinaceae cells

Viability quantification data were analysed for normality. Multifactorial ANOVA was used for examining significance of Symbiodinaceae viability vs. non-viability in *S. viride* faeces, and in contrast to sediment controls. Additionally, the Person Correlation Coefficient was calculated for testing the correlation between healthy and non-healthy Symbiodinaceae populations vs. milligrams of faeces dispersed.

### FCM Method

During the standardization process, three cell populations were gated: P1 (red), P2 (green) and P3 (blue) in a dot plot (“Fig.S2”). Additionally, it was determined an ungated population containing mainly sediment, which hereinafter will be referred to as P4 (black). According to the populations’ distribution, P2 was identified as the one characterized by higher fluorescence signal emission, size and granular cells content. Besides, P3 showed a lower fluorescence signal than P2, and granular cells too; P1 presented the lowest fluorescence signal vs. P2 and P3; and P4 exhibited non-cellular components or debris as it was previously mentioned. Additionally, fluorescence microscopy observations to sorted populations were coherent with the previous light microscopy findings. In this sense, P2 contained Symbiodinaceae cells that presented the highest fluorescence emission; P3 presented zooxanthellae cells with less fluorescence emission in contrast to P2; and finally, P1 and P4 components emitted the lesser fluorescence (“Fig.S4”). Light and fluorescence observations in sorted populations supported the validation of FCM standardized parameters. *S. viride* faeces and control sediment samples were observed under light microscopy to register Symbiodinaceae presence. Zooxanthellae cells were confirmed in 100% of the samples and controls collected. In all cases, cells were observed coccoid in shape with 11-13 um of diameter. Dividing cells stages were evidenced in faeces samples. Intracellular composition patterns varied from well-developed and elongated chloroplasts and defined pyrenoid body to few recognizable organelle’s presence. Additionally, the colour pattern observed corresponded to golden brownish cells to translucent appearance.

### Viability quantification by flow cytometry

Accordingly, to FCM standardized settings, the three gated populations and the ungated population validated were identified and quantified in all *S. viride* faeces and sediment controls (figure 1 in the main text). An entire Symbiodinaceae population per sample was defined as the sum of P2 (viable) and P3 (degrading/unviable) events. Thus, 22855 healthy events were registered in P2 vs. 7500 healthy ones in P3. Regarding faeces or sediment control mg^−1^, a total of 62 healthy events mg^−1^ corresponding to P2 were detected, and in contrast, 32 healthy events mg^−1^ in P3 were evidenced. This fact means a proportion of 2 healthy cells events registered in faeces samples to 1 healthy cell event in sediment control. Conversely, P1 and P4 data were not included in the analyses due to the absence of zooxanthellae in them. In this context, Symbiodinaceae viability calculation per sample was estimated based on the relation between P2 events and the total cells population frequency, expressed as a percentage. The same calculation was done for the estimation of the percentage of degrading/unviable cells (see raw data in “Table S1”).

**Figure 1.**
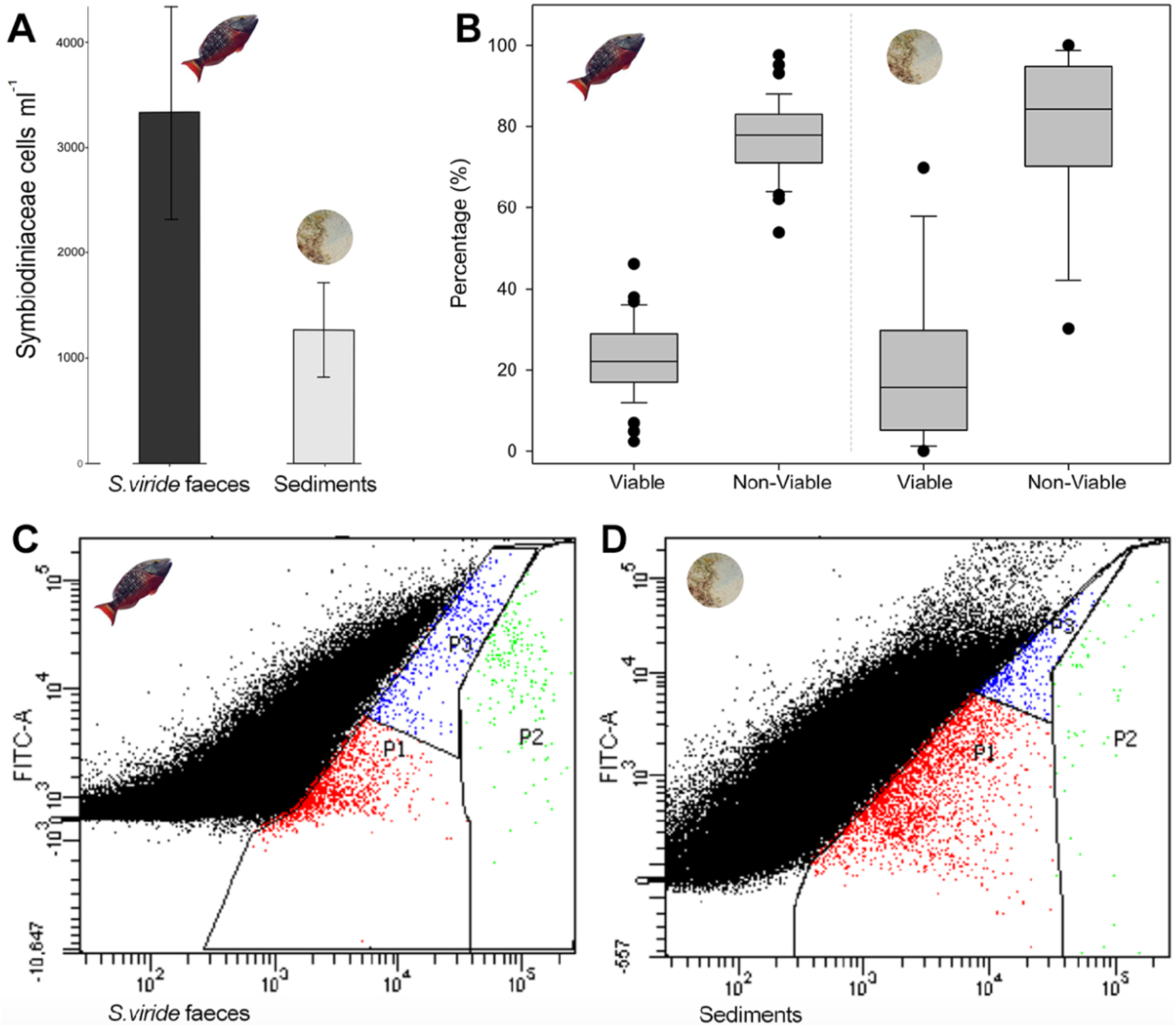
*S. viride* enriches reef sediments with viable Symbiodinaceae cells. Symbiodinaceae enumeration in *S. viride* faeces and reef sediments. We gated two distinctive populations of Symbiodinaceae cells in both faeces and sediments. Viable cells showed an intact cell membrane with spherical nucleus, well developed and elongated chloroplasts, defined pyrenoid bodies and high fluorescence signal. Conversely, degrading/unviable cells exhibited traits such as a fragmented cell membrane, undefined cytoplasmatic material, and a lower fluorescence. A. Cell density in *S. viride* faeces vs. sediment control. B. Boxplot showing significant differences between the percentages of viable cells and degrading/unviable in *S. viride* faeces and sediment controls. No significant differences were observed in the percentage of viable cells in *S. viride* faeces and sediment controls. C-D-Flow cytometry-FCM dot plots displaying the three gated populations identified during standardization in: C. *S. viride* faeces, and D. Sediment controls. Both types of samples, faeces and sediment controls, presented viable cells (high fluorescence) in P2 and degrading/unviable cells in P3.).

### Microbiota in faeces sampling

The microbial component was sampled in West point reef. The sampling was developed in August 2019. Faeces samples were collected in situ through SCUBA diving. Focal *Sparisoma viride* individuals were followed until they defecated. One sample was collected per individual using a 50 mL syringe immediately after the deposition and stored in an individual Ziploc bag. Additionally, sediment control (seabed) samples were collected right next to the faeces deposition. Lastly, three water controls and three sand channel samples were taken aleatorily within the area of study. Water samples were filtered in a 0.2 microns filter. All samples and filters from water samples were transfer to individual testing tubes with ethanol at 70 % to preserve bacteria until DNA extractions in laboratory conditions. A total of 12 faeces samples, 12 sediment samples, three water column samples (controls) and three sand channels samples (controls) were collected.

### DNA extraction and 16S rRNA gene Sequencing

DNA extraction was made using the Quick DNA Fecal/Soil Microbe Kits by Zymo Research according to manufacture instructions. DNA extractions were sent to the Iowa State University (DNA facility) for library preparation and sequencing on the Illumina Miseq Platform. The desired region for amplification was the V4 hypervariable region of the 16S rRNA gene using the primer pairs 515F and 806R(Caporaso et al. 2011).

### 16S rRNA based community analysis

Qiime 2 version 2018.4 was used for bioinformatics analysis with different implemented plugins (Bolyen et al. 2019). Amplicon sequence variants (ASVs) were chosen to work with because they provide better resolution than OTUs based methods(Callahan et al. 2017). First, DADA2 plugin(Callahan et al. 2016) was used for quality control and noise reduction (-p-trim-left-r 0 –p-trim-left-r 0 –p-trunc-len-f 238 –p-trunc-len-r 232). These parameters were chosen according to the reads quality distribution along the length of the sequence. To assess which microbial taxa were present in the samples, we used to q2-feature-classifier plug in to classify our ASVs taxonomically. The reference database used was SILVA 132, specific for 515F and 806R region (http://qiime.org/home_static/dataFiles.html). Next, mitochondria, chloroplast and eukaryotic sequences were removed and a filter of minimum frequency of 17 for the ASVs was applied. These filtered sequences were used to construct the phylogenetic tree, which was made using FastTreeMP with MAFFT alignment implemented in qiime2. The bacterial composition of the environmental controls, water column and sand channels, as well as of sediments and *S. viride* faeces, were graphed using the produced table by the script *qiime taxa collapsed* at family level. Subsequently, the result was graphed using sunburst multilevel pie charts in excel. In order to analyse the diversity of each treatment (*S. viride* faeces, reef sediments, water column and sand bank), we used the package phyloseq (McMurdie and Holmes 2013), implemented in R v 3.5. In addition, we examined the differences between samples richness (Chao1) using ANOVA and Tukey test (“Table S2”). The richness bloxplots were graphed using the package ggplot2(Wickham 2009). For beta diversity a non-metric multidimentional scaling (NMDS) analysis was done using Bray Curtis dissimilarities. A PERMANOVA was performed (999 permutations) in phyloseq to determine differences in the composition of bacterial communities and PERMADISP was performed (999 permutations) to test of multivariate dispersions. Finally, we listed corals and sponges in which *S. viride* usually feeds on, and included the bacterial families associated with those species based on literature data (“Table S3”). This information, along with our findings of the faecal microbiome of *S. viride,* were used to elaborate the Venn diagram found in Figure 3.B.

## Results and discussion

### *S. viride* faeces enriches reef sediments with viable Symbiodinaceae cells

We confirmed the presence of Symbiodinaceae cells in 100 % of *S. viride* faeces and sediment controls, with an average density of 3328 cells ml^−1^ for the former, and 1269 cells ml^−1^ for the latter (See Fig. 1 and Table S1, Fig. S2-4). In general, viable cells in *S. viride* faeces corresponded to 25.97 % ± 1.50, whereas degrading/unviable cells accounted for 74.03 % ± 1.50 (Fig. 1). Similarly, at sediment controls 22.24 % ± 5.36 were viable cells while 77.76 % ± 5.36 corresponded to degrading/unviable cells. The percentages of viable and degrading/unviable cells were significantly different in *S. viride* faeces (F=481.2, *p*<0.001). Contrastingly, no significant differences in the percentage of viable cells were found between *S. viride* faeces and sediment controls (F=0.1, *p*>0.7; Fig. 1). Noticeable, no correlation was observed between the percentages obtained in *S. viride* faeces and control samples vs. dry mass collected (R^2^ =0.023, F=1.067, *p*>0.307), which means that the cell integrity values were not explained by the sample’s mass. Based on these patterns, we propose that *S. viride* may act as a key vector to enrich reef sediments with Symbiodinaceae cells. Therefore, we suggest that *S. viride’s* faeces are crucial for maintaining the main environmental reservoir of dinoflagellates coral symbionts, in accordance to the critical role of sediments for the uptake of symbionts by corals (Adams et al. 2009; Nitschke et al. 2016).

### *S. viride* disperses a highly diverse bacterial community in reef sediments

We analysed 16S rRNA to characterize the bacterial community in *S. viride* faeces and controls. The sediments and feces showed the highest bacterial diversity followed by controls, sand channels and water column samples (Fig. 2a). Alpha diversity index demonstrated that richness of bacterial communities was significantly different between the four types of samples (ANOVA F=9.1, *p*<0.001, Table S2) where sediments and feces present similar richness (p=0.308) and feces differ from controls, the sand channels (p=0.03) and water column (p=0.01) samples. The differences in bacterial composition were assessed using Bray-Curtis dissimilarities in nMDS (Fig. 2b). Particularly, *S. viride* and sediments grouped together and were significantly different from controls (PERMANOVA, F=5.14, R2=0.33, *p*=0.001). Moreover, non-significant differences in the homogeneity of multivariate dispersion were showed between types of samples (PERMADISP, p >0.05).

**Figure 2.**
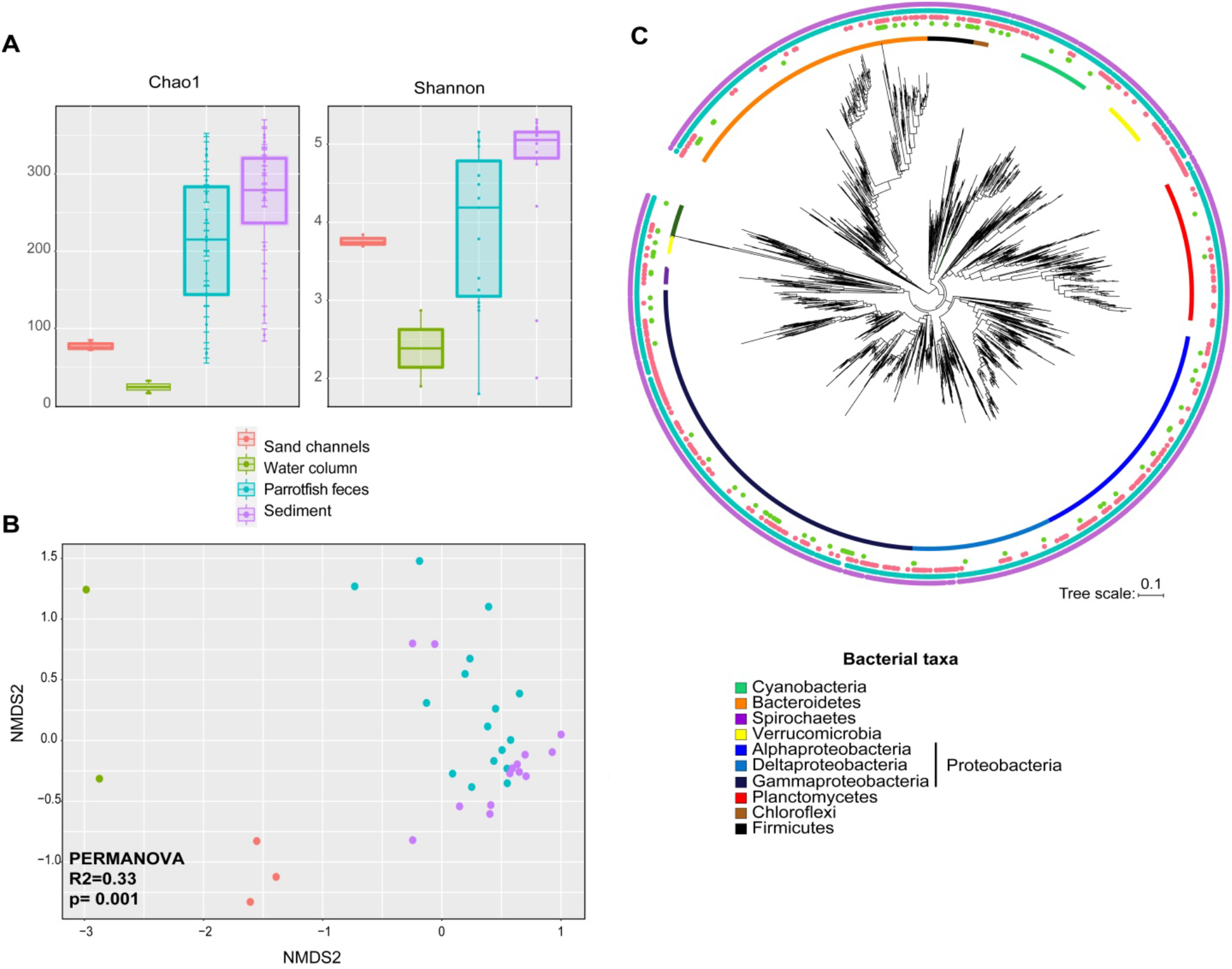
*S. viride* disperses a highly diverse bacterial community in reef sediments. A) Microbial community richness in *S. viride* faeces (blue), sediments (light purple), water column (green) and sand channels (pink) assessed with ANOVA (P-value= 0.003), B) Non metrical multidimensional scaling (nMDS) using Bray Curtis dissimilarities of bacterial communities present in the four types of samples. C) Phylogenetic tree of bacterial community was constructed using maximum likelihood with FasTreeMP and MAFFT alignments. The types of samples are indicated with dots (colours correspond to the part A) above each ASVs tips of the tree. Bacterial phyla are observed as regions above each clade of the tree with different colours. In the case of Proteobacteria, the phylogeny shows the phylum divide in three-class taxonomic levels.

Additionally, Proteobacteria was the most abundant phylum in all the samples with Gammaproteobacteria and Alphaproteobacteria as the dominant classes. This dominance was followed by the phyla Bacteroidetes, Fusobacteria, Firmicutes and Cyanobacteria in both *S. viride* faeces and sediments, and contrastingly, by Firmicutes and Planctomycetes in controls. Regarding the identification of bacterial families of Gammaproteobacteria, *S. viride* faeces were dominated by Vibrionaceae (42.1 %) and followed by Rhodobacteraceae (6.33 %), Fusobacteriaceae (5.53 %), Shewanellaceae (4.48 %) and Woeseiaceae (4.29 %). In contrast, *S. viride* faeces were deficient in Alphaproteobacteria. The Gammaproteobacteria in sediments belonging to Rhodobacteraceae (14.52 %) and Vibrionaceae (14.36 %) were the most abundant families while Halieaceae (6.31 %) and Woeseiaceae (6.06 %) were less represented. We also found the family Cyclobacteriaceae from Bacteroidetes phylum (6.97 %) in the sediments. Although *S. viride* faeces and sediments differed in families abundance, there were similarities in the overall families identified for both types of samples (Figure 3a). Contrarily, the water column (“Fig. S5”) showed higher abundance of bacteria belonging to Phycisphaeraceae (24.07 %), Sphingomonadaceae (17.01 %), Pseudohongiellaceae (13.9 %), Lactobacillaceae (12.86 %), Pseudomonadaceae (11.2 %), while a lesser representation of Vibrionaceae (7.47 %) and, sand channels were dominated by Desulfobulbaceae (11.48 %), Rhodobacteraceae (11.07 %), Sphingomonadaceae (9.41 %) and Arenicellaceae (8.58 %), that differed from faeces and sediments. In fact, it was notable the presence of common amplicon sequence variants (ASVs) across the identified phyla in both faeces and sediments, as it is showed in the phylogeny of the complete bacterial community (Figure 2.c). On this basis, we hypothesize that *S. viride* feaces may support the enrichment of bacteria in reef sediments.

**Figure 3.**
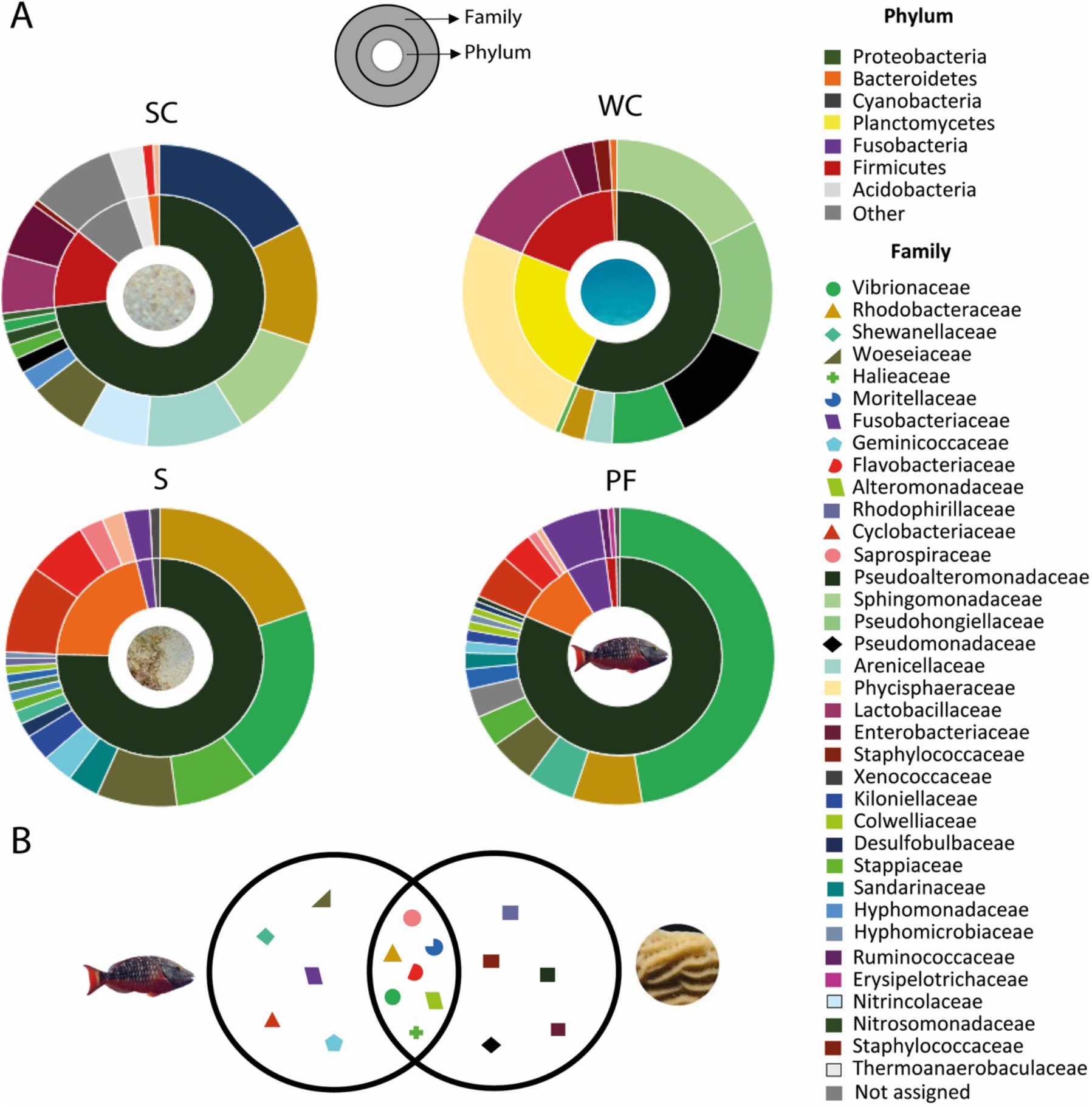
*S. viride* faeces contain a bacterial community commonly associated with the microbiome of foraged species. A) Bacterial composition at phylum and family level using pie charts. Relative abundances of ASVs classified at family and phylum levels are indicated for two types of control samples which are water column (WC) and sand channels (SC); and in the lower part for the microbiota associated with reefs sediments (S) and *S. viride* faeces (PF). B) Venn diagram showing the shared families between *S. viride* faeces (in the left) found in the study and bacterial families of corals (*Agaricia, Orbicella, Siderastrea, Porites*, Diploria and *Pseudodiploria* where parrotfish usually feed) based on literature data (See supplementary table S3) on the right.

### *S. viride* faeces contain a bacterial community commonly associated with the microbiome of foraged species

*S.viride* faeces had high abundances of Gammaproteobacteria ASVs, which are usually found in association with coral tissue (Bourne and Munn 2005). In particular, this bacterial class is dominant in at least ten coral species excavated by *S.viride* (Sanchez et al. 2004; Rotjan and Lewis 2005; Mumby 2009; Castro-Sanguino and Sánchez 2012; Bonaldo et al. 2014; Burkepile et al. 2019b). Eight bacterial families in *S. viride* faeces are associated to coral microbiomes (Fig. 3.b).

Among the Gammaproteobacteria class, the family Rhodobacteraceae, which are prevalent in the microbiome of *Orbicell*a (Meyer et al. 2016) and *Agaricia* (Gonzalez-Zapata et al. 2018), were identified in both *S. viride* faeces and sediments. As diazotrophs, Rhodobacteraceae members are involved in the fixation of nitrogen in association within living coral tissue and the surface mucus layer (Lesser et al. 2018). Similarly, other Gammaproteobacteria metabolise bioproducts from Symbiodinaceae such dimethylsulfoniopropionate (DMSP) and dimethylsufide (DMS) (Raina et al. 2009). As a result, we hypothesize that *S. viride* faeces replenish reef sediments with key bacteria supporting the metabolism of Symbiodinaceae symbionts. This input of Gammaproteobacteria in reefs is relevant in shaping the bacterial community and producing sulphur based antimicrobial functions (Peixoto et al. 2017). Vibrionaceae was the most abundant family identified in *S. viride* faeces, and it is commonly found in the microbiome of several corals and sponges (Dunlap and Pawlik 1996, 1998) (Table S3).

Other families found in *S. viride* faeces were Alteromonadaceae, Flavobacteriaceae and Pseudoalteromonadaceae, which are common bacterial members associated with *Porites* corals (Table S3). Similary, Verrucomicrobiaceae, Vibrionaceae and Rhodobacteraceae are bacterial families present when the mucus is old in *Porites*, probably *S. viride* fed on this coral and disperse through the faeces, however, it have to tested in further studies. On the other hand, *Agaricia* associates closely to Vibrionaceae and Rhodobacteraceae (Gonzalez-Zapata et al. 2018), and *Siderastrea, Porites* and *Pseudodiploria* maintain beneficial relationships with Alteromonadaceae (Table S3) that also are present in parrotfishes of the study, suggest that parrotfish could replenish the coral reef with these symbiont bacteria.

Besides, *Epulopiscium* bacteria was also found in *S. viride* faeces. Although, this result suggests a potential positive feedback of this bacteria to *S. viride* digestion, it should be interpreted with caution in parrotfishes. *Epulopiscium* has a major role in surgeonfish supporting the digestion of marine algae (Ngugi et al. 2017). Parrotfishes are not hindgut fermenters, so the presence of *Epulopiscium* in the *S. viride* could be ingested from sediments and deserves further study (Clements et al. 2014).

### Understanding the ecological feedback of *S. viride’*s microbial dispersal

Our findings suggest that *S. viride* is a vector of symbiotic dinoflagellates, which constantly provides inputs of viable cells. Moreover, a role in replenishing reef sediments is hypothesize based on identifying the same ratio of viable cells:unviable cells in both sediments and faeces. We also found similarities in the bacterial community structure between *S. viride* faeces and reefs sediments. Both of them had diverse bacterial communities and sharing composition. Likewise, our results suggested that the *S. viride* spreads a bacterial community commonly associated to the microbiome of species in which the fish feeds on.

Parrotfish release solid pellets of fine sediment than remain for a short time after being defecated (Goatley and Bellwood 2010). Dispersal occurs in territories of approximately 100 to 300 m^2^ (medium sized), and in a daily feeding range of 50 to 800 m^2^ (Bruggemann et al. 1996). The dispersal of Symbiodinaceae cells and bacteria allows us to start thinking about the resemblance of ecological attributes between *S. viride* and herbivorous in terrestrial ecosystems. The relevance of enriching reef sediments relies on coral health and resilience. Sediments enhance the coral uptake of Symbiodinaceae cells (Adams et al. 2009), which is relevant for coral recruitment and health recovery after bleaching events (Adams et al. 2009; Ali et al. 2019), and facilitate the enrichment of the coral’s ecosphere with beneficial microbes and therefore, supports coral physiological responses from microbial exchange processes (Weber et al. 2019).

## Acknowledgments

Thanks to Nacor Bolaños from Corporación para el Desarrollo Sostenible del Archipiélago de San Andrés, Providencia y Santa Catalina (CORALINA), and Buconos Diving Center team for the assistance during field surveys; Natalia Bolaños from the Laboratorio de Ciencias Básicas Médicas at the Universidad de los Andes supported the FCM analysis process. Humberto Ibarra and his team from the Centro de Microscopía at the Universidad de los Andes contributed with microscopy methods and the members of Laboratorio de Biología Molecular Marina team (BIOMMAR) at the Universidad de los Andes for their valuable assistance.

