## Supplementary Material for "The potential of parrotfish faeces in replenishing reefs with coral-associated microbiome"

**Table S1.** Data from flow cytrometry events (50 ml filtered sample) on *S. viride* faeces including ID_Sample, ID_control, Weigth Total_events, Events P1 ,Events P2, Events P3,Events P4,total_zooxs_p2+p3, Density P2 ml-1, Density P3 ml-1,Density P2+P3 ml-1, percentage_P,2 fluorescence_P2, percentage_P3, fluorescence_P3, fluorescence P1, and Fluoresence P4.

| ID_Sample | ID_Control | Dry_weight | Total_events | Events_P1 | Events_P2 | Events_P3 | Events_P4 | Total_zooxs_P2&P3 | Cell_count_P2 ml-1 | Cell_count_P3 ml-1 | Cell_count_P2&P3 ml-1 | Percentage_Viability_P2 | Fluorescence_P2 | Percentage_Viability_P3 | Fluorescence_P3 | Fluorescence_P1 | Fluorescence_P4 |
|---|---|---|---|---|---|---|---|---|---|---|---|---|---|---|---|---|---|
| M23 | NA | 0,54 | 134984 | 1542 | 54 | 333 | 133055 | 387 | 270 | 1665 | 1935 | 13,95 | 91900 | 86,05 | 22369 | 10165 | 27863 |
| M24 | NA | 0,29 | 83190 | 764 | 44 | 140 | 82242 | 184 | 220 | 700 | 920 | 23,91 | 62634 | 76,09 | 29417 | 12242 | 28914 |
| M25 | NA | 0,22 | 85810 | 1720 | 75 | 262 | 83753 | 337 | 375 | 1310 | 1685 | 22,26 | 82719 | 77,74 | 19757 | 8727 | 25123 |
| M26 | NA | 0,21 | 636944 | 1196 | 101 | 165 | 635482 | 266 | 505 | 825 | 1330 | 37,97 | 113624 | 62,03 | 26891 | 16517 | 42316 |
| M27 | NA | 0,21 | 487296 | 2218 | 9 | 368 | 484701 | 377 | 45 | 1840 | 1885 | 2,39 | 50885 | 97,61 | 16296 | 5402 | 17264 |
| M28 | NA | 0,36 | 126947 | 1495 | 42 | 142 | 125268 | 184 | 210 | 710 | 920 | 22,83 | 75135 | 77,17 | 28806 | 7647 | 27990 |
| M29 | NA | 0,15 | 13865 | 621 | 16 | 100 | 13128 | 116 | 80 | 500 | 580 | 13,79 | 49390 | 86,21 | 27277 | 8820 | 22502 |
| M30 | NA | 0,28 | 86480 | 1399 | 79 | 147 | 84855 | 226 | 395 | 735 | 1130 | 34,96 | 77483 | 65,04 | 30270 | 10598 | 33678 |
| M31 | NA | 0,5 | 5311 | 217 | 4 | 28 | 5062 | 32 | 20 | 140 | 160 | 12,50 | 166370 | 87,50 | 21900 | 7638 | 43689 |
| M32 | NA | 0,23 | 83190 | 1748 | 90 | 326 | 81026 | 416 | 450 | 1630 | 2080 | 21,63 | 73991 | 78,37 | 22771 | 11178 | 25939 |
| M33 | NA | 0,46 | 208116 | 1398 | 62 | 255 | 206401 | 317 | 310 | 1275 | 1585 | 19,56 | 81059 | 80,44 | 36913 | 12915 | 35952 |
| M34 | NA | 0,33 | 73722 | 1036 | 50 | 122 | 72514 | 172 | 250 | 610 | 860 | 29,07 | 83913 | 70,93 | 18563 | 9785 | 30381 |
| M35 | NA | 0,94 | 477661 | 4041 | 23 | 445 | 473152 | 468 | 115 | 2225 | 2340 | 4,91 | 71915 | 95,09 | 16528 | 4964 | 16490 |
| M36 | NA | 0,61 | 1267167 | 4188 | 158 | 436 | 1262385 | 594 | 790 | 2180 | 2970 | 26,60 | 78063 | 73,40 | 24144 | 9274 | 25210 |
| M37 | NA | 0,82 | 1584746 | 6688 | 204 | 812 | 1577042 | 1016 | 1020 | 4060 | 5080 | 20,08 | 76931 | 79,92 | 20827 | 8860 | 22321 |
| M38 | NA | 0,19 | 243744 | 1076 | 51 | 159 | 242458 | 210 | 255 | 795 | 1050 | 24,29 | 86637 | 75,71 | 23364 | 10469 | 29025 |
| M39 | NA | 0,24 | 353440 | 2177 | 43 | 215 | 351005 | 258 | 215 | 1075 | 1290 | 16,67 | 60953 | 83,33 | 22081 | 6454 | 20287 |
| M40 | NA | 0,62 | 94047 | 1129 | 10 | 133 | 92775 | 143 | 50 | 665 | 715 | 6,99 | 73992 | 93,01 | 17735 | 4936 | 21269 |
| M41 | NA | 0,41 | 30389 | 8339 | 4848 | 1613 | 15589 | 6461 | 24240 | 8065 | 32305 | 75,03 | 154340 | 24,97 | 20252 | 95531 | 111155 |
| M42 | NA | 0,44 | 127681 | 1041 | 80 | 222 | 126338 | 302 | 400 | 1110 | 1510 | 26,49 | 97440 | 73,51 | 28361 | 16514 | 38578 |
| M43 | NA | 0,37 | 158165 | 1105 | 48 | 186 | 156826 | 234 | 240 | 930 | 1170 | 20,51 | 97913 | 79,49 | 22718 | 10899 | 29097 |
| M44 | NA | 0,38 | 272741 | 1114 | 112 | 200 | 271315 | 312 | 560 | 1000 | 1560 | 35,90 | 83424 | 64,10 | 25331 | 15888 | 37049 |
| M45 | NA | 0,41 | 788754 | 4388 | 167 | 507 | 783692 | 674 | 835 | 2535 | 3370 | 24,78 | 92094 | 75,22 | 25412 | 8710 | 27203 |
| M46 | NA | 0,48 | 121955 | 2082 | 117 | 426 | 119330 | 543 | 585 | 2130 | 2715 | 21,55 | 78359 | 78,45 | 24197 | 12341 | 28005 |
| M49 | NA | 0,29 | 494957 | 3435 | 103 | 496 | 490923 | 599 | 515 | 2480 | 2995 | 17,20 | 67502 | 82,80 | 23210 | 8434 | 22689 |
| M53 | NA | 0,13 | 102340 | 1200 | 47 | 98 | 100995 | 145 | 235 | 490 | 725 | 32,41 | 90135 | 67,59 | 19979 | 8589 | 26729 |
| M54 | NA | 0,15 | 123610 | 1141 | 34 | 136 | 122299 | 170 | 170 | 680 | 850 | 20,00 | 57470 | 80,00 | 20621 | 7679 | 20362 |
| M55 | NA | 0,41 | 1103542 | 2794 | 270 | 316 | 1100162 | 586 | 1350 | 1580 | 2930 | 46,08 | 97298 | 53,92 | 27076 | 15589 | 41948 |
| M56 | NA | 0,29 | 30389 | 6779 | 4848 | 847 | 17915 | 5695 | 24240 | 4235 | 28475 | 85,13 | 154340 | 14,87 | 24678 | 115534 | 120682 |
| M57 | NA | 0,36 | 357012 | 2440 | 185 | 462 | 353925 | 647 | 925 | 2310 | 3235 | 28,59 | 97637 | 71,41 | 22428 | 13954 | 39503 |
| M58 | NA | 0,18 | 136817 | 1189 | 19 | 103 | 135506 | 122 | 95 | 515 | 610 | 15,57 | 95405 | 84,43 | 24792 | 6560 | 23721 |
| M59 | NA | 0,2 | 166850 | 1135 | 68 | 172 | 165475 | 240 | 340 | 860 | 1200 | 28,33 | 103143 | 71,67 | 23215 | 12992 | 32502 |
| M60 | NA | 0,29 | 57298 | 1162 | 29 | 115 | 55992 | 144 | 145 | 575 | 720 | 20,14 | 60654 | 79,86 | 22162 | 6615 | 21536 |
| M61 | NA | 0,55 | 271671 | 2197 | 105 | 253 | 269116 | 358 | 525 | 1265 | 1790 | 29,33 | 105218 | 70,67 | 22309 | 10728 | 32361 |
| M62 | NA | 0,61 | 216844 | 2849 | 96 | 195 | 213704 | 291 | 480 | 975 | 1455 | 32,99 | 99496 | 67,01 | 22261 | 7764 | 31099 |
| M63 | NA | 0,21 | 1346550 | 2931 | 297 | 511 | 1342811 | 808 | 1485 | 2555 | 4040 | 36,76 | 109914 | 63,24 | 26299 | 18772 | 42868 |
| M64 | NA | 0,41 | 546563 | 2965 | 147 | 567 | 542884 | 714 | 735 | 2835 | 3570 | 20,59 | 100203 | 79,41 | 20252 | 11355 | 32065 |
| M65 | NA | 0,36 | 292011 | 1899 | 187 | 459 | 289466 | 646 | 935 | 2295 | 3230 | 28,95 | 100281 | 71,05 | 26499 | 18618 | 44727 |
| M66 | NA | 0,33 | 66552 | 1492 | 62 | 324 | 64674 | 386 | 310 | 1620 | 1930 | 16,06 | 84380 | 83,94 | 32088 | 13255 | 31188 |
| M67 | NA | 0,45 | 706880 | 2935 | 186 | 661 | 703098 | 847 | 930 | 3305 | 4235 | 21,96 | 102424 | 78,04 | 28168 | 15812 | 34224 |
| NA | CS1 | 0,13 | 4465 | 302 | 14 | 21 | 4128 | 35 | 70 | 105 | 175 | 40,00 | 241366 | 60,00 | 18330 | 15736 | 58590 |
| NA | CS2 | 0,14 | 376 | 7 | 0 | 2 | 3 | 2 | 0 | 10 | 10 | 0,00 | 0 | 100,00 | 12751 | 5912 | 11222 |
| NA | CS3 | 0,13 | 13066 | 348 | 30 | 13 | 99 | 43 | 150 | 65 | 215 | 69,77 | 82268 | 30,23 | 26997 | 12581 | 36737 |
| NA | CS4 | 0,19 | 891 | 25 | 89 | 138 | 639 | 227 | 445 | 690 | 1135 | 39,21 | 91402 | 60,79 | 14619 | 6482 | 29380 |
| NA | CS5 | 0,17 | 222 | 13 | 13 | 55 | 141 | 68 | 65 | 275 | 340 | 19,12 | 104398 | 80,88 | 24300 | 10998 | 36688 |
| NA | CS6 | 0,17 | 2819 | 66 | 351 | 801 | 1601 | 1152 | 1755 | 4005 | 5760 | 30,47 | 74623 | 69,53 | 21224 | 7918 | 20651 |
| NA | CS31 | 0,31 | 17625 | 988 | 4 | 128 | 16505 | 132 | 20 | 640 | 660 | 3,03 | 65510 | 96,97 | 13195 | 4267 | 16032 |
| NA | CS32 | 0,53 | 35626 | 1417 | 48 | 418 | 33743 | 466 | 240 | 2090 | 2330 | 10,30 | 67822 | 89,70 | 18714 | 9845 | 23993 |
| NA | CS33 | 0,26 | 176485 | 4301 | 87 | 722 | 171375 | 809 | 435 | 3610 | 4045 | 10,75 | 64035 | 89,25 | 23398 | 8096 | 21739 |
| NA | CS34 | 0,3 | 73722 | 1036 | 50 | 122 | 72514 | 172 | 250 | 610 | 860 | 29,07 | 83913 | 70,93 | 18563 | 9785 | 26010 |
| NA | CS35 | 0,31 | 6486 | 183 | 2 | 36 | 6265 | 38 | 10 | 180 | 190 | 5,26 | 105849 | 94,74 | 31499 | 9778 | 29427 |
| NA | CS36 | 0,19 | 39480 | 763 | 32 | 141 | 38544 | 173 | 160 | 705 | 865 | 18,50 | 62466 | 81,50 | 23020 | 10286 | 22896 |
| NA | CS55 | 0,21 | 20774 | 1013 | 44 | 100 | 19617 | 144 | 220 | 500 | 720 | 30,56 | 88165 | 69,44 | 13945 | 8293 | 30907 |
| NA | CS66 | 0,3 | 10340 | 440 | 5 | 89 | 9806 | 94 | 25 | 445 | 470 | 5,32 | 58809 | 94,68 | 18030 | 6801 | 20719 |

**Table S2.** Values of the Tukey test for alpha Chao1 richness comparing the four treatments. WC: water column, SC: sand channels, PFF: parrotfish faeces, S: sediments, diff: differences, lwr: lower values, upr: upper values, p adj: *p* value adjusted (bold denotes significant values).

| **Type of sample** | **diff** | **lwr** | **upr** | ***p* adj** |
|---|---|---|---|---|
| WC-SC | -53.000 | -261.013 | 155.013 | 0.899 |
| PFF-SC | 151.933 | 7.817 | 296.049 | 0.035 |
| S-SC | 203.333 | 59.217 | 347.449 | **0.003** |
| PFF-WC | 204.933 | 33.400 | 376.465 | **0.014** |
| S-WC | 256.333 | 84.800 | 427.865 | **0.001** |
| S-PFF | 51.400 | -31.805 | 134.605 | 0.352 |

**Table S3.** Corals where parrotfishes usually feed. Each genus contains the bacterial families found in the different literature references.

| Coral genus | Bacterial Family | Reference |
|---|---|---|
| *Orbicella* | Enterobacteriaceae<br>Vibrionaceae<br>Rhodobacteraceae<br>Flavobacteriaceae, Campylobacteraceae and Burkholderiaceae | [49]<br>[49,50]<br>[28,49]<br>[49] |
| *Agaricia* | Cenarchaeaceae, Burkholderiaceae, Halieaceae, Vibrionaceae,Oxalobacteraceae and Rhodobacteraceae<br>Amoebophilaceae, Rhodospirillaceae, Flammeovirgaceae and Gaiellaceae | [36]<br>[34] |
| *Siderastraea* | Synechococcaceae, Rhodospirillaceae, Nitrosomonadaceae, Desulfurellaceae, Nitrospinaceae, Halieaceae, Saprospiraceae and Rickettsiaceae<br>Corynebacteriaceae, Bacillaceae<br>Endozoicomonaceae<br>Aquabacteriaceae, Comamonadaceae, Enterobacteriaceae, Flavobacteriaceae, Lachnospiraceae, Micrococcaceae, Phyllobacteriaceae, Planctomycetaceae, Pseudoalteromonadaceae, Pseudomonadaceae, Rhizobiaceae, Rhodobacteraceae, Rikenellaceae, Ruminococcaceae, Staphylococcaceae, Streptococcaceae and Ulvophyceae<br>Rhodospirillaceae<br>Alteromonadaceae | [51]<br>[52,53]<br>[52]<br>[53]<br>[51,53]<br>[50] |
| *Porites* | Rhodobacteraceae<br>Alteromonadaceae<br>Synechococcophycideae<br>Endozoicomonaceae<br>Verrucomicrobiaceae<br>Vibrionaceae | [34,54]<br>[54]<br>[55]<br>[34,55,56]<br>[34]<br>[28,34,50] |
| *Diploria* | Endozoicomonaceae<br>Vibrionaceae, Microccocaceae | [55]<br>[57] |
| *Pseudodiploria* | Halomonadaceae, Moritellaceae and Alteromonadaceae<br>Pirellulaceae, Rhodopirallaceae and Flavobacteriaceae | [28]<br>[58] |

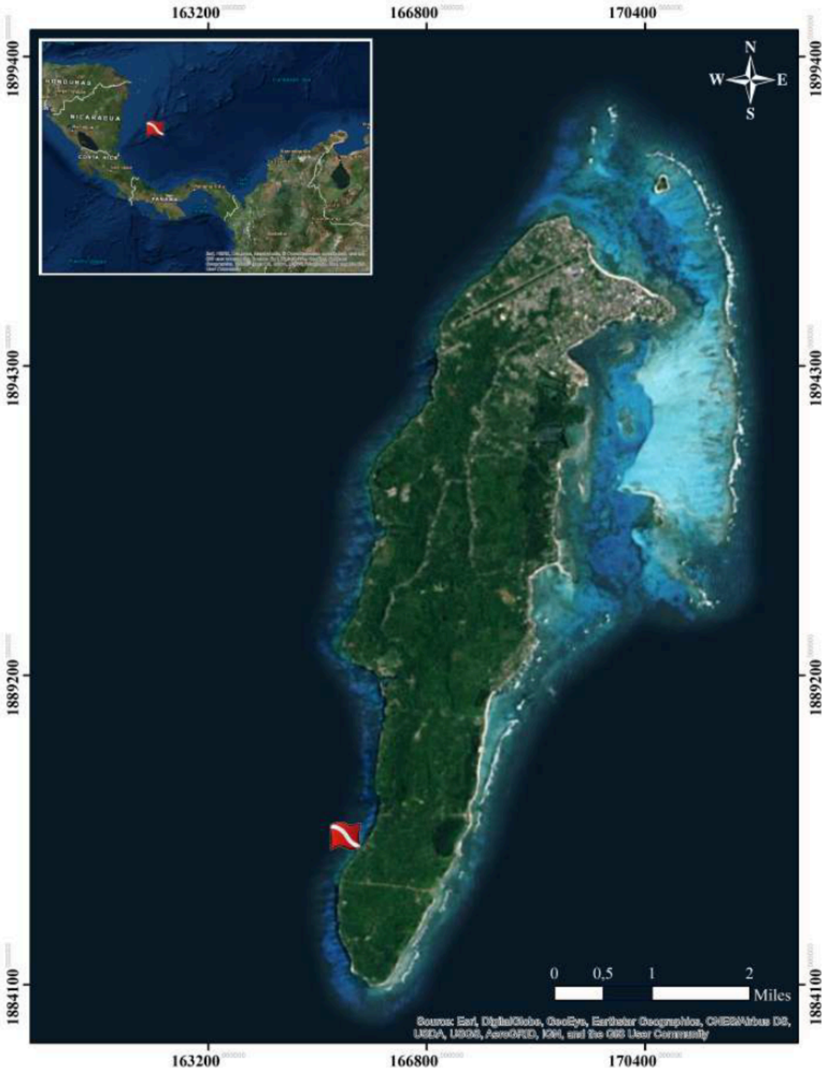


**Fig. S1.** Map of San Andrés island, Southwestern Caribbean, showing the sampling site.

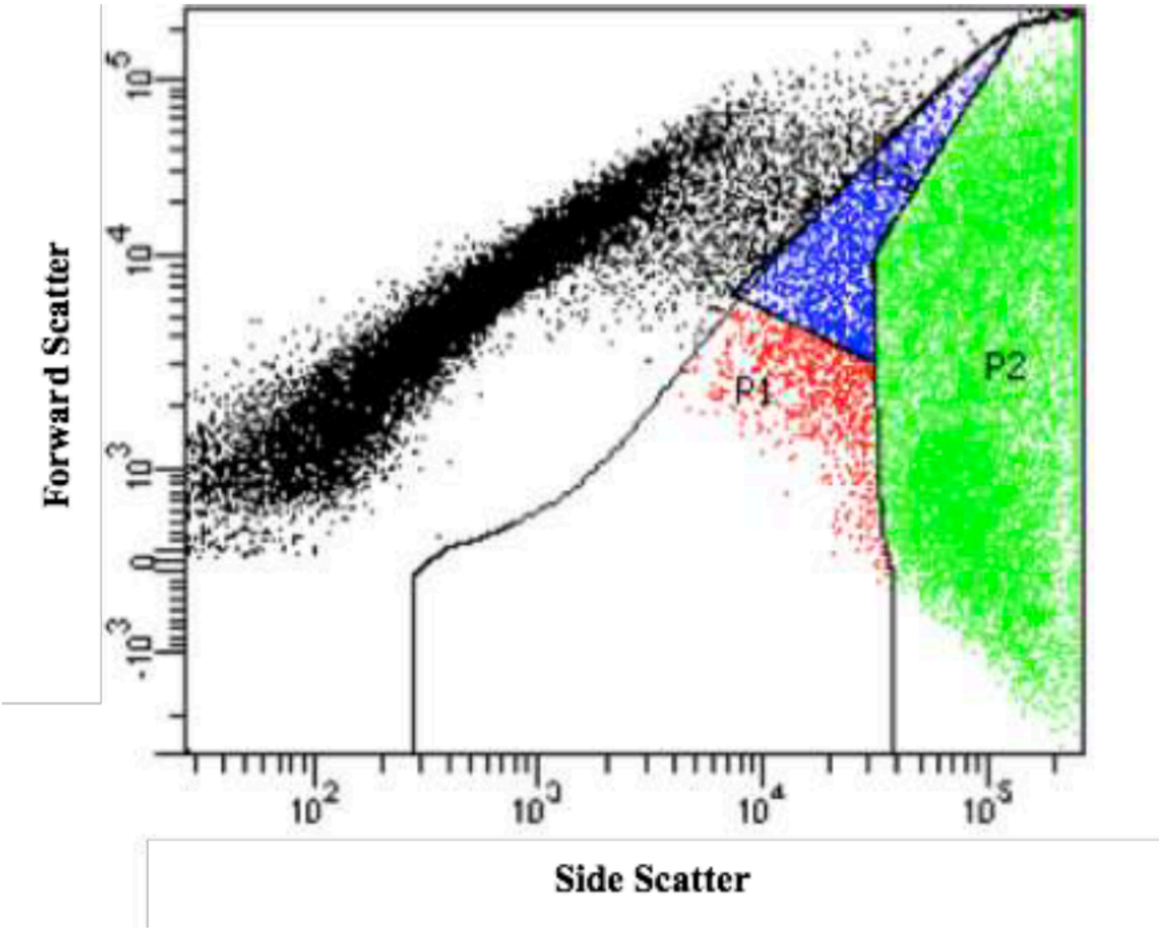


**Fig. S2.** Cell populations identified during FCM standardization process. Complementary, sorting results conduced to validate the three gated populations P1 (red), P2 (green) and P3 (blue) . An ungated population containing mainly sediment was called P4. Specifically, viable/high fluorescent *Symbiodinaceae* cells were found in the P2 population, and degrading/unviable *Symbiodinaceae* cells were registered in P3. On the other hand, none *Symbiodinaceae* cells were identified in P1, in which macroalgae fragments and unspecific subproducts were observed in accordance with *S. viride* diet.

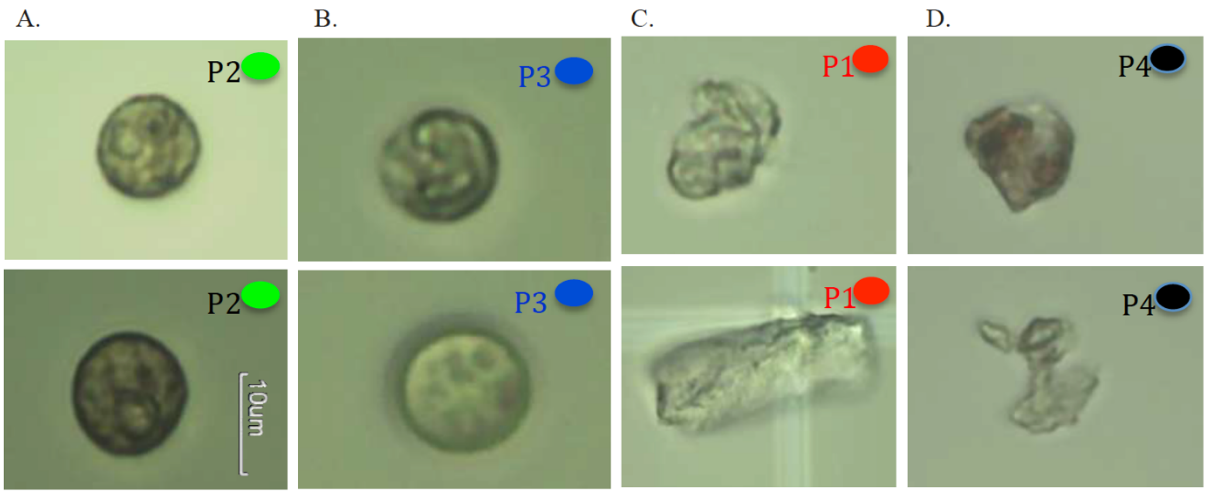


**Fig.S3** Sorted population identities validation: A. P2 contains *Symbiodinaceae* with viable/high fluorescent appearance; B. P3 presents *Symbiodinaceae* with degrading/unviable characteristics; C. P1 as well as D. P4 does not contained *Symbiodinaceae*-like cells.

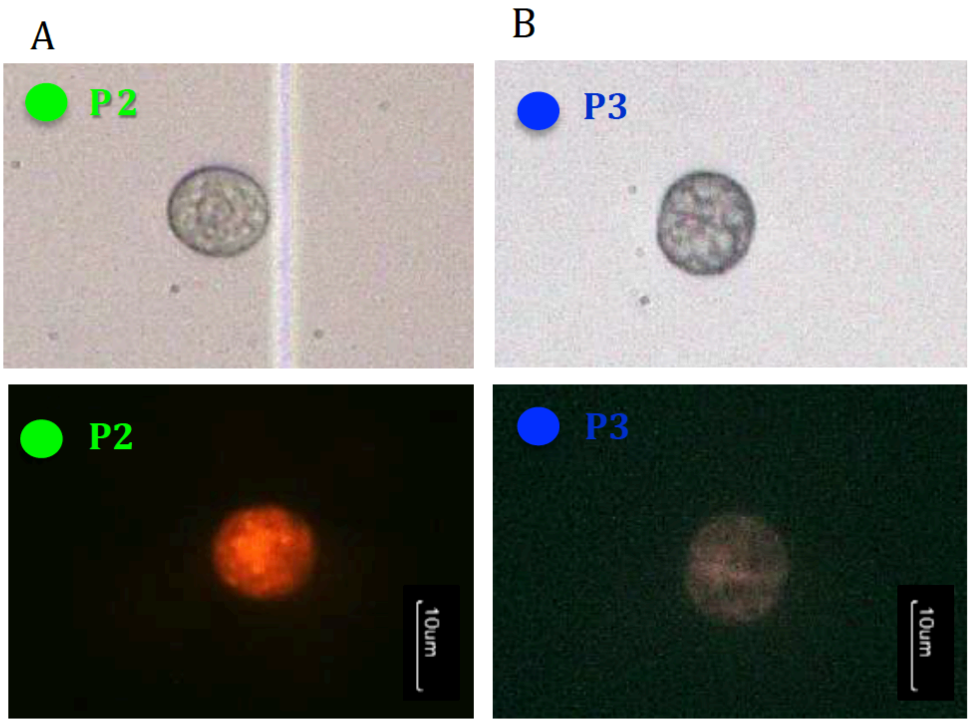


**Fig.S4** Light vs. fluorescence microscopy registers to sorted cells populations (40x): A. P2 *Symbiodinaceae* cell emitted higher fluorescence than P3. B. P3 *Symbiodinaceae* cell with its characteristic fluorescence.
